# A new insight into RecA filament regulation by RecX from the analysis of conformation-specific interactions

**DOI:** 10.1101/2022.03.14.484239

**Authors:** Aleksandr Alekseev, Georgii Pobegalov, Natalia Morozova, Alexey Vedyaykin, Galina Cherevatenko, Alexander Yakimov, Dmitry Baitin, Mikhail Khodorkovskii

## Abstract

RecA protein mediates homologous recombination repair in bacteria through assembly of long helical filaments on single-stranded DNA (ssDNA) in an ATP dependent manner. RecX, an important negative regulator of RecA, is known to inhibit RecA activity by stimulating the disassembly of RecA nucleoprotein filaments. Here we use a single-molecule approach to address the regulation of (*E. coli*) RecA-ssDNA filaments by RecX (*E. coli*) within the framework of distinct conformational states of RecA-ssDNA filament. Our findings revealed that RecX effectively binds the inactive conformation of RecA-ssDNA filaments and slows down the transition to the active state. Results of this work provide new mechanistic insights into the RecX-RecA interactions and highlight the importance of conformational transitions of RecA filaments as an additional level of regulation of its biological activity.

## Introduction

In bacteria, the RecA protein is a central player in DNA repair, genetic recombination and SOS response activation [1–5]. RecA acts through assembly of long helical filaments on single-stranded DNA (ssDNA) in an ATP-dependent manner [6–10]. RecA-ssDNA filaments can further pair with homologous duplex DNA to form a D-loop, assisting in the rescue of stalled replication forks [11,12]. RecA also catalyses a DNA strand exchange reaction, a key step during homologous recombination which ensures faithful DNA repair and genetic recombination. Apart from that, RecA-ssDNA filaments stimulate proteolytic cleavage of the LexA repressor, allowing activation of over 40 SOS-response genes involved in DNA repair and cell cycle regulation [13,14]. Importantly, RecA-dependent SOS-response activation is one of pathways for development of antibiotic resistance by bacteria through enhanced mutagenesis in the presence of continuous DNA damage. RecA nucleoprotein filaments are very dynamic, and may experience large-scale conformational changes induced by ATP binding and hydrolysis [6,15–18]. Besides intrinsic ATPase activity of RecA, assembly and stability of RecA-ssDNA filaments are also dynamically regulated by a network of various protein mediators [19–23].

RecX is an important negative regulator, which has been reported to suppress ATPase, DNA-pairing, and strand-exchange activities of RecA [24–28]. RecX also inhibits RecA co-protease activity [29]. Genes encoding RecX were found in genomes of a wide diversity of bacteria and some plants [30]. In *Escherichia coli*, recX is a SOS-regulated gene located downstream of recA which encodes a small 19.4 kDa protein. recX and recA are co-transcribed, however expression of recX is down-regulated at both transcriptional and translational levels resulting in about a 500-fold lower protein level compared to RecA [31].

*In vivo* studies showed that either loss-of-function RecX mutation or overexpression of RecX decrease bacterial resistance to UV irradiation [29], while overexpression of RecA resulted in deleterious effects when RecX was mutated [32–35]. For some species, such as *Deinococcus radiodurans*, RecX shows dual negative regulation of RecA function - it not only directly inhibits RecA activity at the protein level but also inhibits RecA induction at the transcriptional level [36]. Interestingly, RecX is also an important mediator of natural transformation in *Bacillus subtilis*, where it colocalizes with RecA threads, while the lack of RecX decreases chromosomal transformation approximately by 200-fold [27].

A capping model has been proposed to elegantly explain the inhibitory mechanism of RecX at substoichiometric concentrations relative to RecA. According to this model, RecX binds to the growing 3’-end of the RecA filament and blocks the filament extension which results in net filament depolymerisation [37]. On the other hand, observation of faster RecA depolymerization at higher RecX concentrations together with structural evidence of RecX binding along the RecA filament groove [24,25,38] indicate that alternative internal-nicking mechanism may exist, in which RecX locally destabilizes the RecA filament and increases the number of the disassembling ends [28,39]. In support of this model, a recent study of *Mycobacterium smegmatis* RecX (MsRecX) showed that incubation of MsRecX with RecA filaments resulted in RecA dissociation from within the filament. It has also been shown that mechanical forces, which are an important factor in regulating the stability of the RecA filament [26,40], can counteract the inhibitory effect of RecX at as little as 7 pN, prevent disassembly of the RecA filament, and even stimulate the repolymerization of RecA on DNA in the presence of RecX.

Another completely different model of the RecX inhibitory action was proposed based on the transmission electron microscopy data [25]. This model is not associated with a decrease in the number of RecA monomers on DNA as a result of filament depolymerization, but alternatively suggests that RecX binds to the interface between RecA monomers and blocks the transition between inactive and active states, thereby preventing ATP hydrolysis.

Single-molecule studies proved to be a powerful tool for addressing the inhibitory mechanism of RecX. Magnetic tweezers assays have been used to directly observe disassembly of RecA filaments induced by RecX proteins from *Mycobacterium tuberculosis* [41] and *Bacillus subtilis* [26]. Measurements of the dynamic characteristics of RecA-ssDNA filaments in the active state showed that RecX promotes net depolymerization of preformed filaments at low tensile forces (about 3 pN) in ATP hydrolysis and RecX concentration dependent manner, which is generally consistent with results of ensemble experiments. At the same time, this approach made it possible to discover that depolymerization of the RecA filament took place in a non-monotonic stepwise manner with pauses of various lengths (10-100 s). Moreover, RecA depolymerization could proceed with an initial lag phase followed by a strikingly rapid net depolymerization phase [26] indicating a complicated nature of interaction between RecX and RecA-DNA complex.

Recently, we characterized three mechanically distinct conformational states that occur within the *E. coli* RecA-ssDNA filaments in the course of ATP hydrolysis [18]. In the present work, building on the developed single-molecule assay we aimed to study interactions of *E. coli* RecX with different forms of RecA nucleoprotein filaments to further deepen understanding of RecX inhibitory mechanism.

## RESULTS

### RecX stimulates biphasic shortening of RecA-ssDNA filaments

In this work we addressed the dynamics of interaction between *E. coli* RecX and RecA nucleoprotein filaments (Figure 1A) utilizing a single-molecule approach which combines dual-trap optical tweezers and a microfluidic laminar flow cell for manipulation of individual DNA molecules (Figure 1B) as reported elsewhere [18,42–48]. The microfluidic flow cell (Lumicks B.V.) consisted of 5 laminar flow channels which facilitated optical trapping of the beads, DNA tethering and RecA nucleoprotein filaments assembly (see Methods). Additionally, changing the composition and order of the microfluidic channels provided flexibility in the experimental design to study how RecX interacts with various forms of RecA-DNA filaments.

**Figure 1.**
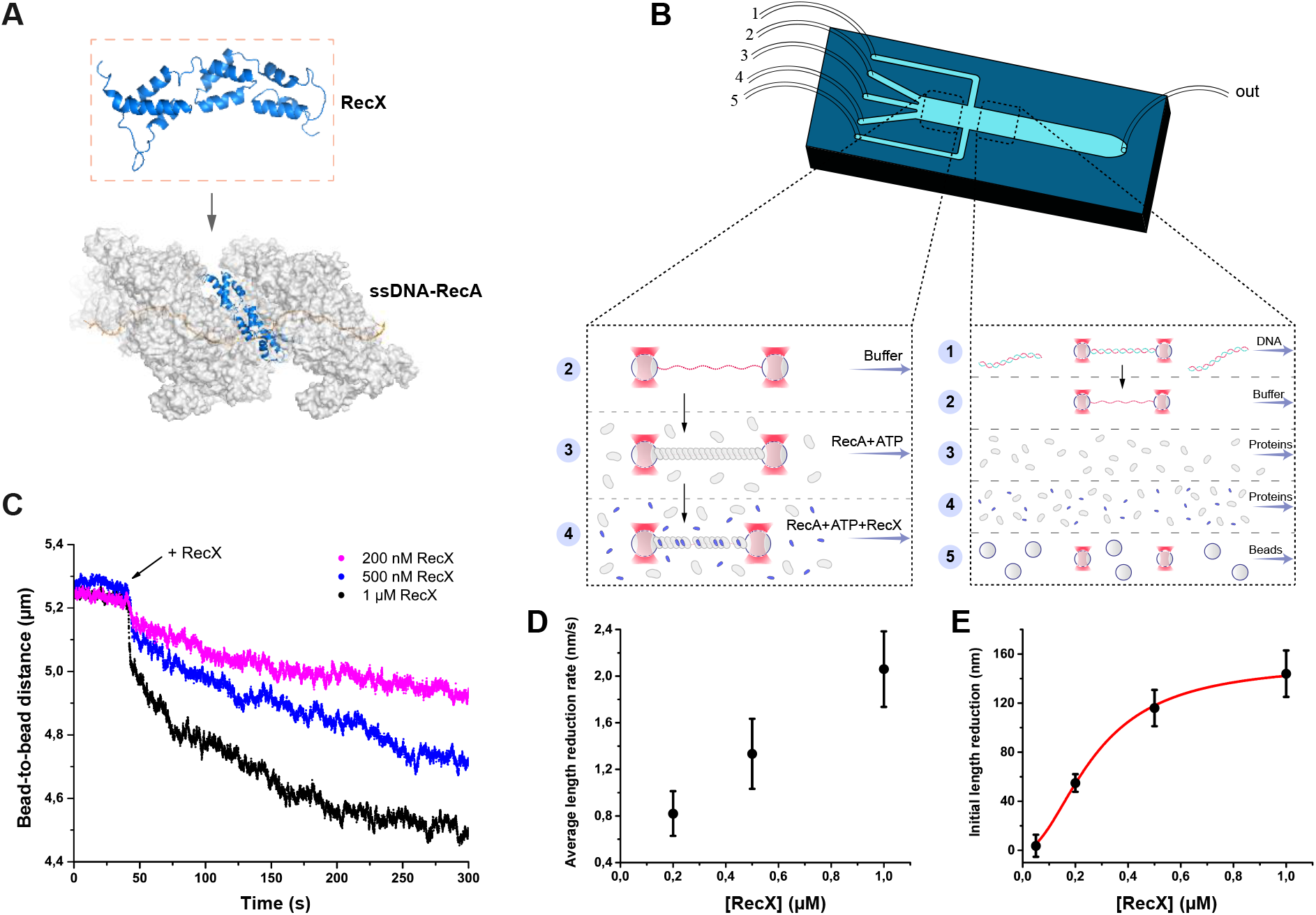
The study of the RecX effect on the RecA-ssDNA filaments. (a) A schematic of RecX binding along the groove of the active RecA-ssDNA. Atomic structure model for RecA::RecX::ssDNA is adopted from [38]. (b) A schematic of a 5-channel microfluidic flow cell (Lumicks). Dash line highlights two working regions. The three-channel region was used to study the effect of RecX on the RecA-ssDNA filament. In the 5-channel region the beads trapping, DNA-tether formation and generation of ssDNA by force-induced melting were performed. (c) The change in the length of RecA-ssDNA filament upon transition from the channel containing 1 μM RecA and 1 mM ATP to the channel containing 1 μM RecA, 1 mM ATP and various concentrations of RecX. During incubation a constant tension of 3 pN was applied to the tether. (d) The impact of RecX concentration on the average rate of reduction in the RecA-ssDNA filament length over 250 seconds after initial steep decrease. (e) The dependence of the RecX induced initial sharp decrease in RecA-ssDNA filament on the RecX concentration. Solid curve - fit of experimental data with Hill equation with a Hill coefficient of 2.0 ± 0.3. Each data point in (d) and (e) is a mean value of at least three measurements, bars represent standard deviation. **Figure 1-source data 1** Source data for traces of RecA-ssDNA filaments, average length reduction rate and initial length reduction values.

Previously, single-molecule magnetic tweezers experiments revealed that *M. tuberculosis (Mt)* RecX and *B. subtilis (Bs)* RecX stimulate gradual disassembly of the corresponding active RecA-ssDNA filaments resulting in the net decrease of the DNA tether length [26,41]. First, we aimed to investigate whether *E. coli* RecX (further referred as RecX) exhibits the same behaviour. To address the dynamics of RecA filaments preassembled on 11,071 nt long single-stranded DNA (further referred as RecA-ssDNA filaments), the RecA-ssDNA tether was stretched with a constant force of 3 pN while its end-to-end distance was simultaneously recorded. The force was chosen as low enough to not introduce structural deformations in the filament as reported previously [18].

Introduction of the preassembled RecA-ssDNA filament into the channel containing RecX (0.2 – 1 μM) in the presence of both RecA (1 μM) and ATP (1 mM) resulted in the initial step-like shortening of the tether followed by a slow decrease in the end-to-end distance (Figure 1C). After 200 seconds of incubation with RecX the length reduction reached steady state and proceeded with an almost linear profile. Average rate of the filament length reduction was dependent on the concentration of RecX (Figure 1D). Higher RecX concentrations stimulated faster shortening in line with previously reported observations for *Mt* and *Bs* RecX [26,41]. Interestingly, the initial step-like shortening of the RecA-ssDNA filament also scaled with RecX concentration (Figure 1E). Corresponding dependence is presented in figure 1E and is well fitted by a Hill equation with a Hill coefficient of 2.0 ± 0.3. These observations indicate that *E. coli* RecX stimulates shortening of the RecA-ssDNA filaments which proceeds in two phases.

#### Shortening of RecA-ssDNA filaments induced by RecX is driven by reversible conformational change in the filament and slow filament disassembly

At a stretching force of 3 pN the length of the RecA-ssDNA filament is significantly greater than the length of bare ssDNA [6,18] (Figure S1). Therefore, shortening of the RecA-ssDNA tether can be caused by RecX actively displacing RecA from ssDNA. Alternatively, since both RecX and RecA were present at comparable concentrations, RecX could interact with free RecA in solution preventing its interaction with DNA.

Additionally, RecA-ssDNA filaments may adopt different conformations depending on the nucleotide cofactor bound in the RecA monomer-monomer interface. ATP-bound or “active” form is characterized by a longer pitch which results in a stretched conformation of the filament [49–55], while “inactive” ADP-bound and *apo* forms are characterized by a smaller pitch and a more compact structure [49–53,56,57]. Hence, shortening of the RecA-ssDNA tether induced by RecX can be caused not only by RecA dissociation, but also by a conformational transition between active and inactive states.

To further elucidate the details of the RecX interaction with RecA nucleoprotein filaments we performed experiments without the presence of free RecA during the incubation with RecX. For this, we first preassembled the RecA-ssDNA filament in the channel containing RecA and ATP and transferred the tether into a channel containing only the buffer supplemented with ATP (Figure 2A). Upon transfer, RecA-ssDNA tether did not show any significant change in its length and was relatively stable. This confirmed that RecA is tightly bound to ssDNA and no turnover between DNA-bound and free RecA is necessary to keep RecA-ssDNA filaments stable.

**Figure2.**
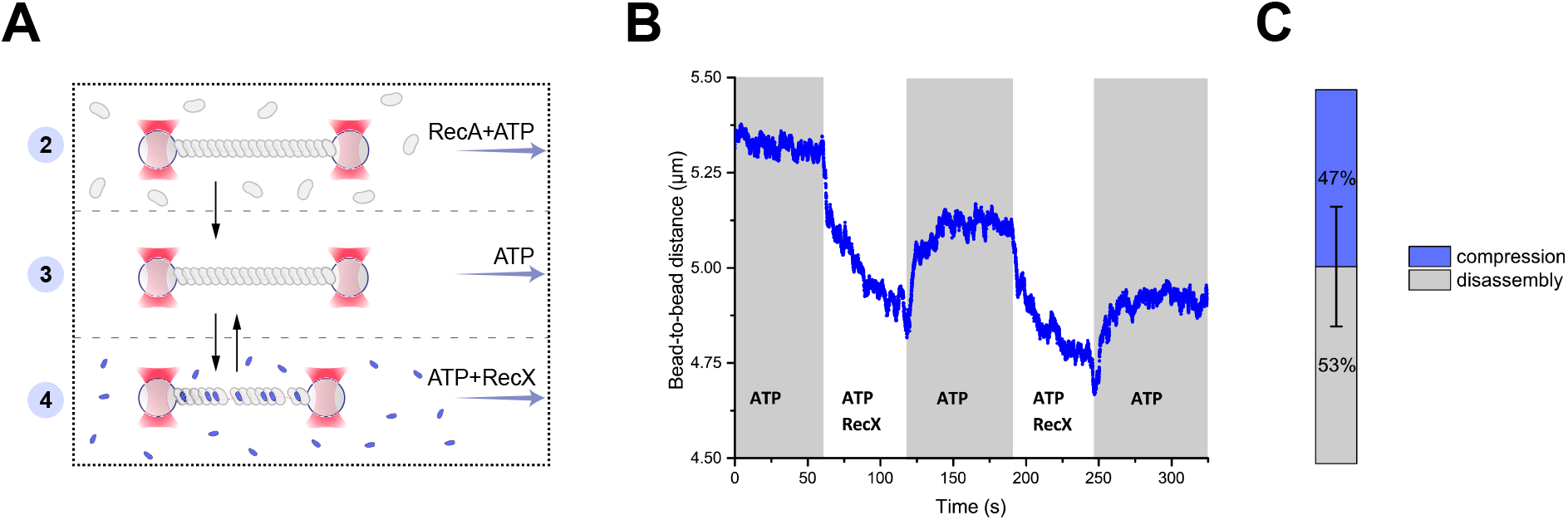
RecX induced reversible changes in the RecA-ssDNA filament structure. (a) A schematic of the experiment revealing that RecX is able to induce reversible structural changes in the RecA-ssDNA filaments. (b) RecX induces reversible changes in the RecA-ssDNA filament structure in the presence of ATP. (c) A comparison of the reversible (compression) and the irreversible (disassembly) reduction in RecA-ssDNA filament length. Stacked histogram represents multiple measurements for six different molecules. Bars represent standard deviation. **Figure 2-source data 1** Source data for RecX-induced reversible changes in the length of RecA-ssDNA filament.

Next, the RecA-ssDNA filament was transferred into a channel containing ATP (1 mM) and RecX (1 μM). This transition resulted in the reduction of the tether length (Figure 2B) in a manner similar to the experiment described in the previous section, indicating that shortening of the RecA-ssDNA filament is induced by direct interaction of RecX with the DNA-bound RecA since this process does not depend on the presence of free RecA. Surprisingly, when the RecA-ssDNA filament was moved back to the channel containing ATP without RecX a gradual elongation of the tether length was observed (Figure 2B). Since no free RecA was present, the observed elongation is solely driven by structural rearrangement within the RecA-ssDNA filament. This indicates that binding of RecX to the RecA-ssDNA tether introduces a conformational change within the RecA nucleoprotein filament that leads to a shortening of its overall length which can be reversed when RecX is eliminated.

Cycles of shortening and elongation of the tether could be observed multiple times on the same DNA molecule by simply transferring the RecA-ssDNA filament between the RecX and the buffer channels (Figure 2B). However, after each incubation with RecX subsequent length restoration was not complete with the RecA-ssDNA filaments consistently showing decreased length. This indicates that at each cycle a part of RecA irreversibly dissociated from DNA (Figure 2C). These observations suggest that RecX-induced shortening of the RecA-ssDNA tether is a combined result of two processes: a fast conformational transition into a more compact form and slow disassembly of the RecA nucleoprotein filament.

#### RecX interacts with inactive RecA-ssDNA filaments and inhibits transition into the active state

At saturated concentrations of ATP (1mM used in the current work) the RecA-ssDNA filament adopts an overall “active” or stretched conformation. However, continuous ATP hydrolysis occurs throughout the filament and generates a dynamic and heterogeneous structure in which *apo* and ADP conformations may transiently occur as local patches [18,47,58]. One possible explanation of the reversible shortening of the RecA-ssDNA tether is that RecX interacts with the compressed conformations locally occurring within the filament and increases their lifetime. To test this hypothesis, we addressed the interaction of RecX with the *apo* form of RecA-ssDNA filaments.

We first assembled an active filament in the presence of 1 μM RecA and 1 mM ATP and then transferred it into a channel containing a buffer without ATP (Figure 3A). ATP hydrolysis stimulated fast accumulation of the compact *apo* conformation accompanied by a characteristic reduction of the tether length (Figure 3B). Next, the filament was transferred into the channel containing 500 nM RecX and incubated for 100 s. During incubation no change in the filament length was registered. After that, the filament was transferred back into the buffer channel and subsequently into the RecA-ATP channel.

**Figure 3.**
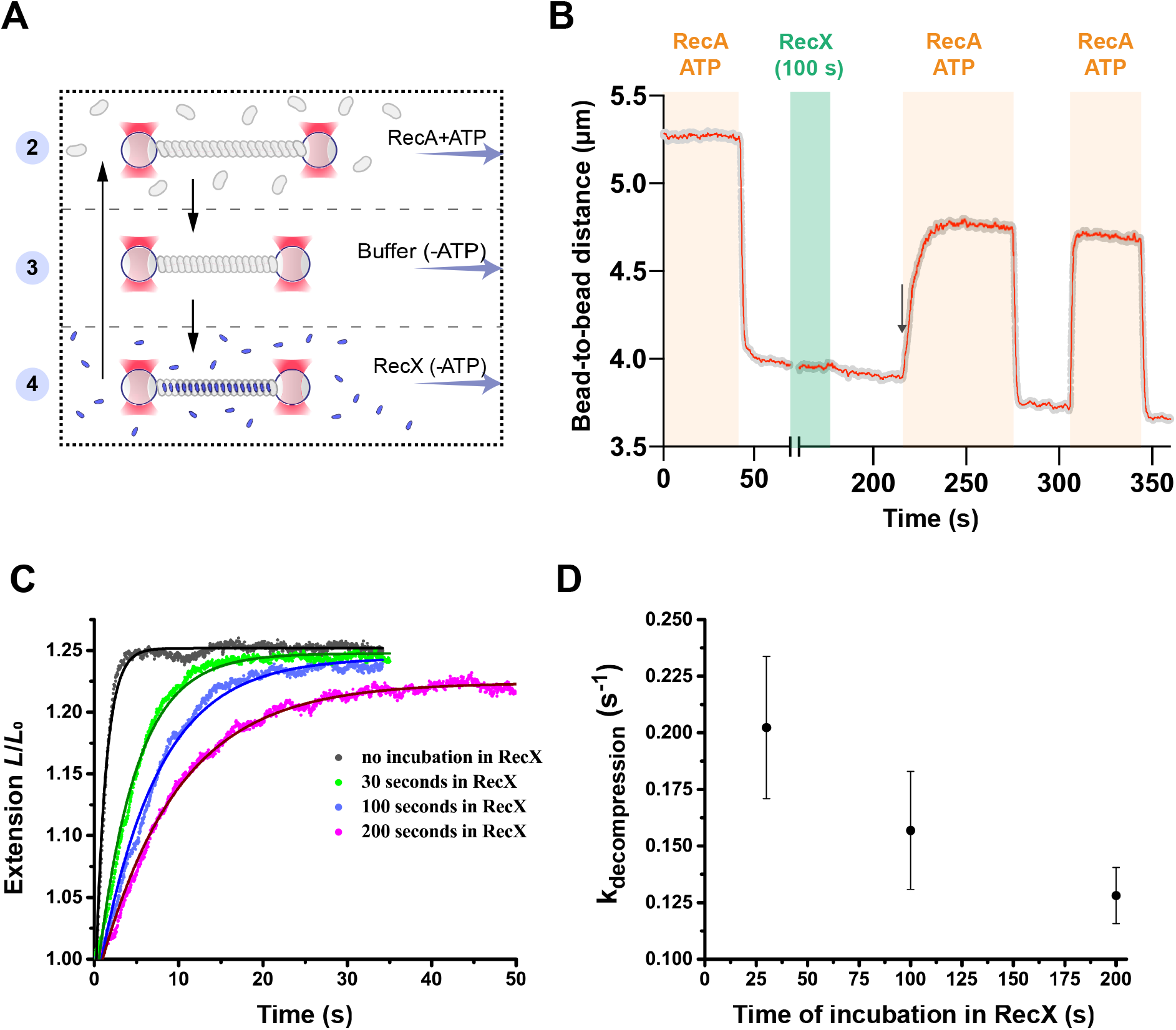
RecX affects the conformational transition of RecA-ssDNA filament from the inactive state to the active. (a) A schematic of the experiment revealing that RecX binds inactive RecA-ssDNA filaments. (b) The change of the RecA-ssDNA filament length upon conformational transitions between *apo* and ATP-bound states. Incubation of *apo* RecA-ssDNA filament with 500 nM RecX (green area) leads to a slowdown of the subsequent decompression of the RecA-ssDNA filament (black arrow points the beginning of the slowed down decompression). A constant tension of 3 pN was applied to the tether during incubation and transitions. (c) Relative extension of the RecA-ssDNA filament in the course of decompression after incubation of inactive RecA-ssDNA filament with 500 nM RecX for 30, 100 and 200 seconds. (d) Corresponding rate constants of the decompression obtained by exponential fitting (solid line in (c)) of the elongation profiles. Each point is a mean of at least six measurements. Bars represent standard deviation. **Figure 3-source data 1** Source data for slowdown decompression of apo RecA-ssDNA filament caused by incubation with RecX.

In the RecA-ATP channel we observed a slow elongation of the tether suggesting gradual accumulation of the stretched ATP conformation within the RecA-ssDNA filament due to a conformational transition from the *apo* into the ATP-bound state. However, decompression of the filament after incubation with RecX was remarkably slower than when RecX incubation was omitted (Figure 3C). Interestingly, when the same molecule was subjected to another cycle of *apo*-ATP transition without RecX incubation, decompression proceeded rapidly with the rate similar to the control case without RecX incubation. This experiment confirms that RecX interacts with the *apo* form of the RecA-ssDNA and delays its conformational transition into the ATP-bound form. However, decompression of the RecA-ssDNA filament irreversibly disrupts a specific linkage between RecX and the *apo* state of the filament.

To quantitatively assess the observed effect we varied the time of incubation with RecX and calculated the rate constants of the corresponding decompression events using exponential fitting (Figure 3C,D). A profile of the filament elongation was fitted with the following expression: *L*=*L_0_+ΔL·exp^(-kt)^*, where *L_0_* is the length of the *apo* state of the RecA-ssDNA filament, *ΔL* - the change of RecA filament length upon conformational transition from the *apo* to the ATP-bound state, *k* - rate constant of the conformational transition. Increasing the time of incubation with RecX from 30 to 200 seconds resulted in slowing down of the decompression characterized by decrease in k_decompression_ from 0.2 to 0.13 s^-1^. For comparison, the rate constant of the RecA-ssDNA filament decompression without incubation with RecX was 1.3 ± 0.3 s^-1^ (N = 5). This value is limited by the temporal resolution of the current measurement, e.g. the time required to transfer the tether between the channels while operating optical tweezers in a force clamp mode. In the previous work using the same DNA construct and a fast movement between microfluidic channels the rate constant of the *apo*-ATP transition was estimated to be greater than 10 s^-1^ [18].

Additional control experiments verified that RecX remains bound to the *apo* form of the RecA-ssDNA filament in the RecX-free buffer (Figure S2). The observed slowdown in the decompression rate was also independent of how long the RecA-ssDNA filament remained in the inactive state in the absence of RecX (Figure S3). Our results indicate that RecX tightly binds the *apo* state of RecA-ssDNA filament and stabilizes it, retarding the *apo*-ATP transition. The conformational transition of the RecA-ssDNA filament from the *apo* to the ATP state results in the gradual loss of specific interaction of RecX and the *apo* conformation. Similar results were obtained for the interaction of the RecX with the ADP-bound state of RecA-ssDNA filament (data not shown).

#### ATP promotes RecX dissociation from RecA-ssDNA filaments

To understand whether RecX remains bound to the RecA-ssDNA filament after ATP-induced decompression we generated a fluorescent version of *E. coli* RecX. For this, the N-terminus of RecX was fused to a green fluorescent protein mNeonGreen [59,60] via a short peptide linker. mNeonGreen-RecX (RecX_mNG_) retained the inhibitory effect on RecA at the level comparable to the wild-type RecX. This was tested by ability of RecX_mNG_ to inhibit ATPase activity of RecA in bulk (data not shown) as well as to slow down the *apo*-ATP transition rate in the single-molecule experiments (Figure 4A).

**Figure 4.**
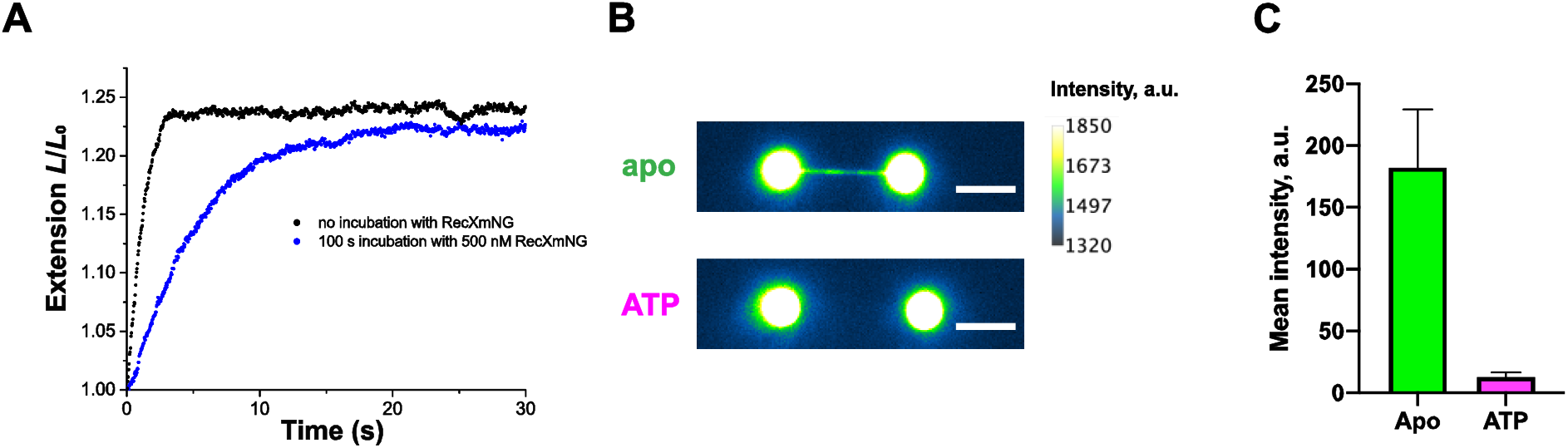
Fluorescent visualization reveals that RecX dissociates from the ATP-bound state of the RecA-ssDNA. (a) Relative extension of the RecA-ssDNA filament in the course of apo-ATP transition without incubation in RecX_mNG_ (black curve) and after incubation of apo RecA-ssDNA filament with 500 nM RecX_mNG_ for 100 seconds (blue curve). (b) Fluorescent images of RecA-ssDNA filament in apo and ATP-bound state after incubation with 1 μM RecX_mNG_ for 30 seconds. Scale bar is 5 μm. (c) Comparison of the average intensity of the tether for apo (N = 6 molecules) and ATP-bound RecA-ssDNA filament (N = 3 molecules) after incubation with RecX_mNG_ (consistently with b). Data are representative of three independent experiments and values are expressed in mean ± SD. **Figure 4-source data 1** Source data for RecX_mNG_-induced slowdown decompression of RecA-ssDNA filament and average intensity values for apo and ATP-bound RecA-ssDNA filaments after incubation with RecX_mNG_. **Figure 4-source data 2** Raw tiff images of apo and ATP-bound RecA-ssDNA filaments after incubation with RecX_mNG_.

To evaluate binding of RecX_mNG_ to the *apo* form of RecA-ssDNA filaments we first preassembled the active RecA-ssDNA filament and transferred it into the channel with ATP-free buffer to obtain the *apo* conformation. After that, the tether was incubated in the channel containing 1 μM of RecX_mNG_ for 30 seconds and then transferred back to the buffer channel to be imaged using wide-field fluorescence microscopy. Visualization revealed a strong fluorescent signal along the DNA tether confirming that RecX_mNG_ tightly binds the compressed RecA-ssDNA filament (Figure 4B). For fluorescence experiments a longer DNA substrate of ~24,000 nt was used to obtain a larger separation between the beads.

Next, we transferred the RecX_mNG_-bound filament to the channel containing RecA and ATP to induce a transition into the active state and imaged the tether again after decompression was completed. Fluorescent visualization demonstrated significant loss of RecX_mNG_ from the RecA-ssDNA filament (Figure 4B). Quantitative analysis showed that the conformational transition of RecA filaments to the ATP-bound state results in reduction of the average intensity of RecX_mNG_ bound to the tether almost down to a background level (Figure 4C).

To address dynamics of the *apo*-ATP transition we performed continuous fluorescent visualisation of the RecA-ssDNA filament during the transfer from the ATP-free to the ATP containing channel (video 1). To keep a constant stretching force during image acquisition one of the beads was released from the optical trap and the tether was stretched by a constant fluid flow adjusted to exert a force of ~3 pN. Upon entry into the ATP containing channel the dynamic increase in the tether length was correlated to the gradual loss of the RecX_mNG_ fluorescence signal. These observations suggest that ATP-induced transition between inactive and active states of the RecA filament promotes dissociation of RecX.

### Video 1

#### RecX efficiently disassembles RecA filaments on double-stranded DNA

In addition to RecA filaments assembled on ssDNA we decided to test whether RecX exhibits similar effects on RecA filaments formed on double-stranded DNA (dsDNA). Since, the assembly of RecA-dsDNA filament is impeded at room temperature, the experiment was carried out at 37 °C. RecA-dsDNA filaments were assembled by applying a tension of 55 pN to the dsDNA molecule (11,071 bp) in the presence of 1 μM RecA and 1 mM ATP (Figure 5A). As a result of RecA binding the length of the dsDNA molecule increased to 5.4 ± 0.1 μm (N = 9) which corresponds to ~150% elongation of the B-form of dsDNA. At a force of 3 pN RecA-dsDNA filaments exhibited the length of 4.9 ± 0.2 (N = 9), which is slightly lower than the length of RecA filaments formed on equivalent ssDNA at the same force, however this length was stable over a long period of time.

**Figure 5.**
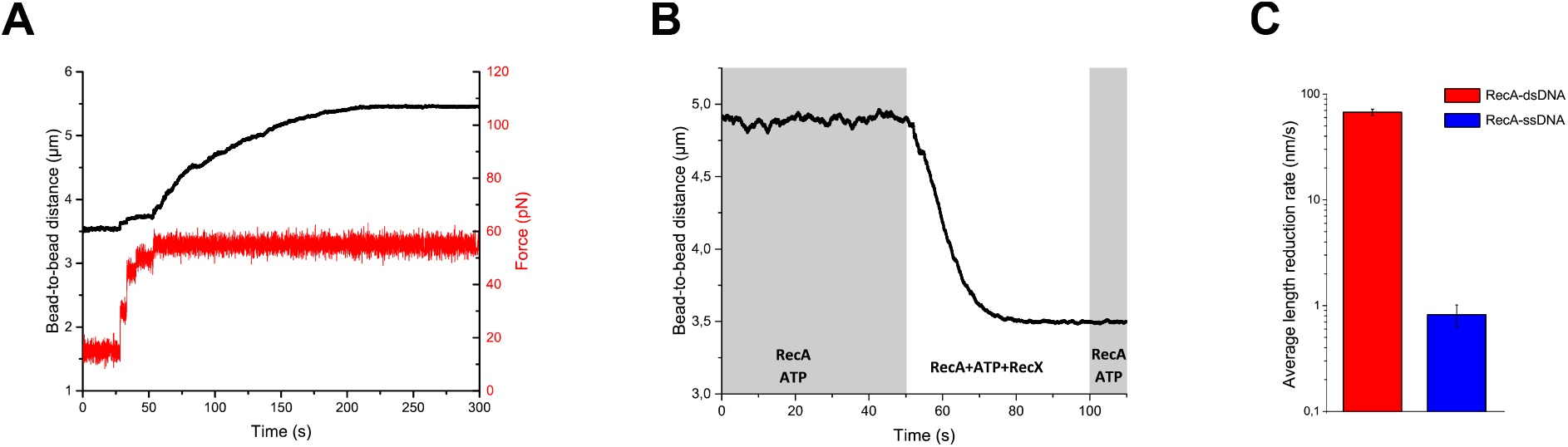
RecX effectively promotes disassembly of RecA-dsDNA filaments. (a) The assembly of the RecA-dsDNA filament. (b) The disassembly of the RecA-dsDNA filament in the presence of 200 nM RecX. (c) The comparison of the average length reduction of RecA-dsDNA (N = 4) and RecA-ssDNA (N = 4) filament induced by 200 nM RecX. The data for RecA-ssDNA is consistent with figure 1D. Data are representative of at least three independent experiments and values are expressed in mean ± SD. **Figure 5-source data 1** Source data for RecA-dsDNA filament assembly profile, RecX-induced disassembly of RecA-dsDNA filament and average length reduction rate for RecA-dsDNA and RecA-ssDNA filaments in the presence of RecX.

Preassembled RecA-dsDNA filaments were transferred from the channel containing 1 μM RecA and 1 mM ATP to the channel containing 1 μM RecA, 1 mM ATP and 200 nM RecX (Figure 5B) at a constant tension of 3 pN. Surprisingly, in this case RecX stimulated a dramatic reduction of the tether length down to the length of bare dsDNA within 30 seconds. Subsequent reintroduction of the tether back to the RecX-free channel did not produce any elongation of the dsDNA molecule. These observations suggest that RecX is highly efficient at irreversible disassembly of RecA-dsDNA filament. Analysis of the results obtained with 200 nM RecX revealed that the average rate of the filament length reduction is almost 2-order higher for the RecA-dsDNA filaments compared to RecA-ssDNA (Figure 5C).

## Discussion

The results presented above comprise a detailed analysis of the conformation-specific interactions of the *E. coli* RecX and RecA nucleoprotein filaments. We first discuss the effect of RecX on the RecA-ssDNA filaments. Our experiments confirm that RecX stimulates a net disassembly of the RecA-ssDNA filament, however this process is very slow and takes hundreds of seconds for a ~5 micrometer long RecA filament to depolymerise. Previously proposed capping model [37] suggests that RecX binds to the filament end and blocks its growth leading to a net RecA depolymerisation. This model alone fails to explain our results since in the absence of RecA polymerisation RecX actively stimulated shortening of the RecA-ssDNA filament (Figure 2). Such behaviour indicates that RecX also binds along the RecA-ssDNA filament and promotes structural rearrangements and/or disassembly from within the filament.

Unexpectedly, we discovered that RecX interacts with the compressed *apo* form of the RecA-ssDNA filament and inhibits its transition into the ATP-bound state. This is in line with the reported inhibition of the RecA ATPase activity by RecX [39] and directly shows that RecX can effectively block ATP hydrolysis by preventing the corresponding conformational transitions within the RecA-ssDNA filament without actual displacement of RecA monomers from ssDNA as proposed in [25]. It is worth to note that RecX-bound inactive RecA filaments persisted without noticeable changes for over 200 seconds indicating that RecX does not promote depolymerisation of inactive RecA filaments.

Interestingly, when such filaments were supplemented with ATP, RecX gradually dissociated in the course of apo-ATP transition (Figure 4). Previously, low resolution electron microscopy demonstrated that RecX can bind along the groove of the ATP-bound conformation of RecA filaments [25], however these interactions were addressed at a higher concentration of RecX (3 μM). Our results indicate that RecX strongly binds the *apo* conformation of the RecA-ssDNA filament while its affinity to the ATP-bound conformation is significantly lower. Depolymerisation of the RecA-ssDNA filament occurred only in the presence of ATP (Figure 2B), suggesting that RecX-stimulated RecA dissociation from DNA is coupled to ATP binding or hydrolysis.

Based on the observations discussed above we propose a following model of how RecX interacts with RecA-ssDNA filaments under conditions of continuous ATP hydrolysis (Figure 6).

**Figure 6.**
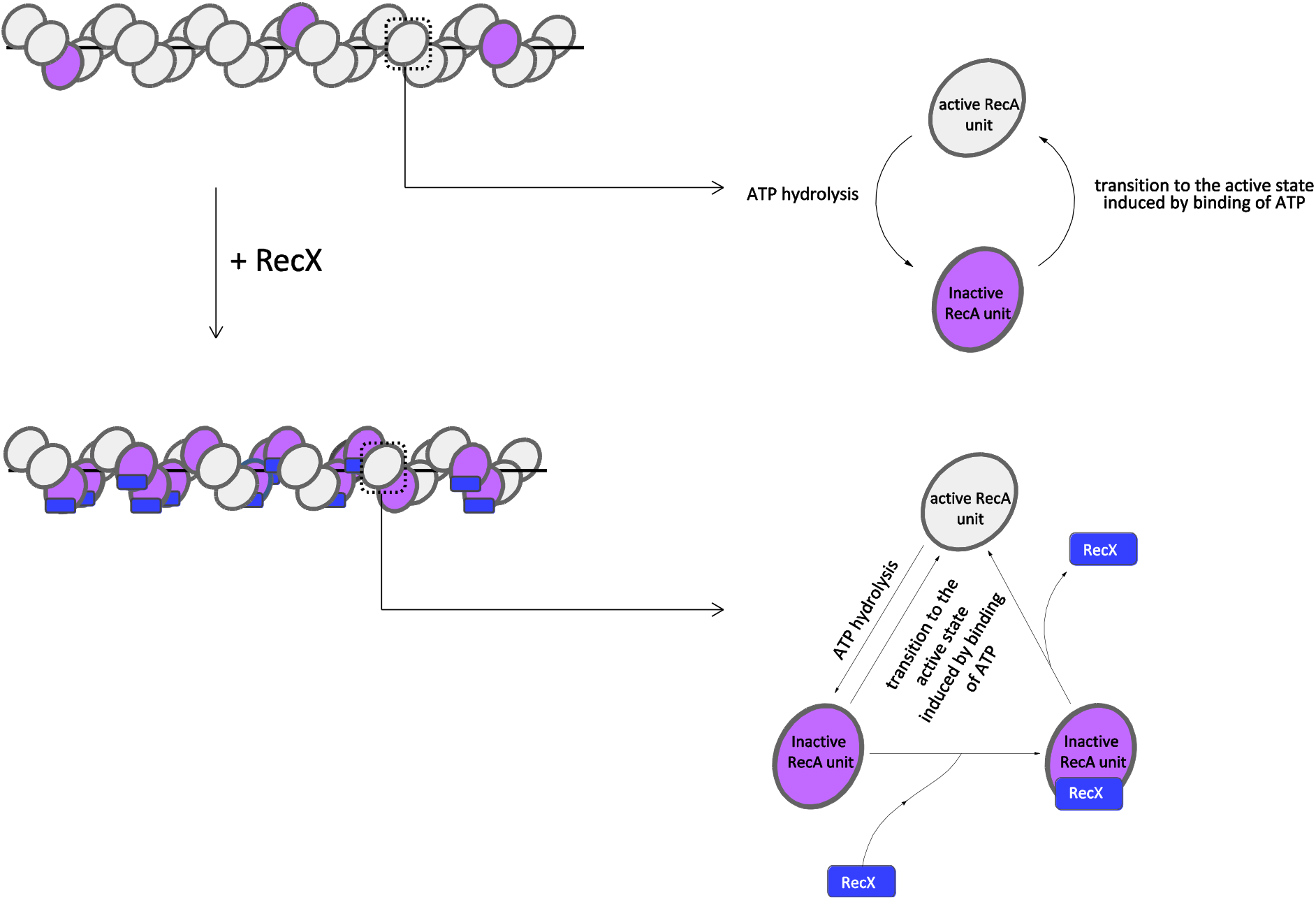
Model of RecX interaction with RecA-ssDNA filaments under conditions of continuous ATP hydrolysis. In the presence of ATP RecX binds inactive patches within RecA-ssDNA filaments and hampers the transition to the active state (see text for details).

RecX binds the inactive patches, transiently presented within the active RecA-ssDNA filament, and increases their lifetime by hampering the transition into the active state. The transition of the inactive patch bound by RecX to the active state is accompanied by dissociation of RecX. As a result of the dynamic interaction of RecX with inactive states, their average lifetime and fraction within the filament increase, which leads to the reduction in the filament length. When RecX is removed from solution, bound RecX dissociates as the inactive patches change their conformation to the active, the fraction of inactive states returns to the initial level.

The current study showed that RecX-induced fast compression of the RecA-ssDNA tether is characterized by positive cooperativity with the Hill coefficient ≈2 (Figure 1E). However, such fast compression is in contrast to the slow binding of RecX to the compressed RecA filament which apparently does not reach saturation even after 100-seconds incubation with a micromolar concentration of RecX (Figure 3C). Previous studies revealed that the apo-ATP transition of the RecA-ssDNA flament is highly cooperative [15,18] meaning that such transitions within the filament are affected by the conformational state of adjacent RecA monomer-monomer interfaces. We hypothesize that apart from inhibition of conformational transitions by direct binding, RecX may interfere with the cooperativity of the apo-ATP transition in the process that involves binding of two RecX monomers.

Unlike RecA-ssDNA complexes, RecX is much more efficient at disrupting RecA filaments formed on duplex DNA. Such filaments mimic postsynaptic RecA-DNA complexes, yet their structure is closely related to the presynaptic RecA-ssDNA filament [55]. Considering structural similarity, we assume that RecX interacts with the RecA-dsDNA filament in a similar manner by trapping transiently occurring inactive states. However, the internal tension of the stretched duplex DNA makes the compressed form of the RecA filament highly unstable. RecX impedes the transition into the stable active conformation and by this stimulates the dissociation of RecA (Figure 5B). Instability of the inactive form of RecA-dsDNA filaments was confirmed by a control experiment which demonstrated that elimination of ATP leads to a complete dissociation of RecA from dsDNA within seconds (Figure S4A). Alternatively, RecA-dsDNA filaments depolymerisation may be due to efficient inhibition of the filament growth by RecX via capping mechanism [37]. RecA-dsDNA complexes are not stable without free RecA and quickly depolymerize as a result of ATP hydrolysis, indicating that a significant turnover of free and DNA-bound RecA is necessary for maintaining the filament (Figure S4B).

One potential implication of high efficiency of RecX-induced disruption of RecA-dsDNA filaments is preventing erroneous binding of RecA to a duplex DNA inside the cell. Another interesting possibility is that RecX may interact with the RecA-ssDNA filament and stay bound until the homologous dsDNA is found. Once the strand exchange occurs and the heteroduplex is formed within the filament, RecX stimulates fast dissociation of the postsynaptic complex.

Results of this work provide new mechanistic insights into the RecX-RecA interactions and highlight the importance of conformational transitions of RecA filaments as an additional level of regulation of its biological activity. Regulation of recombination by conformation-specific interactions might be relevant for eukaryotic recombinases Rad51, Dmc1.

## Materials and methods

### DNA constructs and proteins

A linear 11.1 kbp DNA with biotinylated ends was used for all single-molecule manipulations except fluorescence experiments. This DNA construct was prepared as described previously [18,47]. Briefly, the biotin labelling was performed by a ligation of the oligonucleotides 5’-XXXXXCAGTCCAGCT-3’ and 5’-CTAGCGAGTGXXXXX-3’, where X is biotin tag, to the plasmid vector prl574 containing insertion of the rpoC gene [61] digested with XbaI and SacI restriction enzymes. Single-stranded DNA molecules were generated during experiment by a force-induced melting of double-stranded DNA molecules [62].

For fluorescence experiments a longer DNA construct was used. A set of 22 kbp and 24 kbp DNA molecules, enabling generation of ssDNA by force induced melting, was prepared starting from Lambda DNA (48.5 kbp) as follows. The 3’-ends of Lambda DNA were filled in with biotinylated nucleotides. The reaction contained 1X Tango Buffer (Thermo Scientific), 50 μM dATP, dTTP, dGTP (Thermo Scientific), Biotin-16-dCTP (Jena Bioscience), 7.5 nM Lambda DNA (New England Biolabs), 0.3 units/μl Klenow fragment (Thermo Scientific). After incubation at 37°C for 30 min the reaction was heat inactivated (10 min at 75°C), afterwhich DNA was purified using Bio-Gel P-30 size-exclusion spin column (Bio-Rad). Then DNA was digested with SacI (Thermo Scientific) in 1X Tango Buffer and subsequently ligated with the biotinylated oligonucleotide 5’-XXXXXCAGTCCAGCT-3’ at 22°C for 2 hours. A 50:1 ratio of oligonucleotides to DNA overhangs was used. Short complementary oligonucleotide 5’-GGACTG-3’ was added to increase ligation efficiency [63]. The reaction was heat inactivated (20 min at 65°C), after which DNA was purified from the excess of oligonucleotides using the Bio-Gel P-30 spin column.

Wild type EcRecA and EcRecX were purified as described previously [37,64,65]. To visualize RecX interaction with RecA-ssDNA filament, a fluorescent RecX-mNeonGreen (RecX_mNG_) fusion protein was prepared as follows. A genetic construct consisting of 6xHis-tag, short linker (GMASM), mNeonGreen gene, short linker (GGGSGG), and recX (*E. coli)* was cloned into the pBAD/HisB plasmid vector. *E. coli* Rosetta cells were transformed with the resulting pBad_mNeonGreen_RecX vector. Bacteria cells were grown at 37°C. When OD600 reached 0.6, expression was induced by addition of 1 mM isopropyl 1-thio-D-galactopyranoside (IPTG), after which cells were grown for 3 hours. The cells were harvested by centrifugation and resuspended in a lysis buffer (Na-phosphate buffer, pH 7.5, 500 mM NaCl, 5% glycerol, 10 mM Imidazole, 1 mg/ml Lysozyme). Cells were sonicated for 30 minutes and were centrifuged for 40 minutes at 16,000g. After passing through a 0.4 μm filter the supernatant was loaded onto a HisTrap HP 1 mL column (GE Healthcare). The protein was eluted with a lysis buffer supplemented with 300 mM Imidazole. The eluted fractions were loaded onto a Superose 6 Increase 10/300 GL column (GE Healthcare) equilibrated with 50 mM Tris–HCl pH 8.0 (4°C), 500 mM NaCl, 1 mM DTT. The resulting chromatogram revealed a single peak in the 50 kDa region, corresponding to the monomeric fraction of the mNeonGreen-RecX fusion protein (47.6 kDa). The fractions containing RecX_mNG_ were supplemented with 10% glycerol, frozen in liquid nitrogen and stored at −80°C. The resulting RecX_mNG_ protein showed the ability to inhibit the ATPase activity of RecA at a similar level compared to the wild-type RecX protein.

### Optical tweezers setup

Custom-built dual optical tweezers were used as described previously [18,66]. In brief, optical trapping was performed with the Nd:YVO4 1064 nm CW laser (5 W, Spectra-Physics BL-106C), high numerical aperture oil immersion lens (LOMO, 100X, NA = 1.25), 5-channel microfluidic flow chip (Lumicks), EMCCD camera (Andor Technology, iXon Ultra 897). The x,y-position of one of the traps was additionally controlled with nanometre accuracy by the mirror mounted on a piezo platform (S-330.80L, Physik Instrumente, Karlsruhe, Germany). The applied force and the end-to-end distance of the DNA tether were measured in real time with 30-ms time resolution by processing images of the trapped beads with a custom-made software designed in LabVIEW. The optical trap stiffness was calibrated by a drag force method using a high-precision piezo stage (P-561.3DD, Physik Instrumente). To apply a constant tension to DNA tether, optical tweezers were operated in a force clamp mode.

Fluorescent images of RecX_mNG_-RecA-ssDNA complexes were obtained with a separate EMCCD camera (Cascade II, Photometrics) using freely available MicroManager software. The fluorescence was excited by the DPSS 473 nm CW laser (100 mW, Lasever, LSR473U) attenuated with a 2.0 OD ND filter. Filter set 10 (Carl Zeiss) was used for separating the fluorescence and the excitation light. Images were collected at 50 ms exposure time and EM gain of 3000 and saved as TIFF files without compression. Images were further processed in FIJI ImageJ.

### Single molecule assay

#### Buffer solution, experimental conditions, and passivation of the microfluidic chip

All experiments were made in 25 mM Tris-HCl (pH 7.5), 5 mM MgCl_2_, 50 mM NaCl. ATP containing channels were also supplied with 10 U/ml pyruvate kinase and 0.2 mM phosphoenolpyruvate. In fluorescence experiments, microfluidic channels used for imaging were also supplemented with the oxygen scavenging system: 3 u/ml pyranose oxidase (Sigma-Aldrich), 90 units/ml catalase (Sigma-Aldrich), 1% glucose. Channels of the microfluidic chip were passivated with 0.5 % Pluronic F-127 and 0.1 % BSA [46]. Experiments with RecA-ssDNA filaments as well as experiments studying DNA binding activity of RecX were done at 22C°. Experiments with RecA-dsDNA filaments were made at 37C°.

#### DNA-tether formation and force-induced melting

Two of the five channels of the microfluidic chip (Figure S5A) were correspondingly fed with a 0.01% solution of a 2.1-μm streptavidin-coated polystyrene beads (Spherotech) and a 15 pM solution of dsDNA (11.1 kbp) with biotinylated ends (Figure S5E). To obtain a single DNA tether, two beads were initially optically trapped and then moved to the DNA containing channel, where attachment of the DNA to the beads occured. The attachment of a single DNA molecule to the beads was verified by applying a stretching force of about 20 pN to the tether and controlling its length. Additional check was done during force-induced melting, when a double-stranded DNA molecule exhibited an overstretching plateau at a tension of about 65 pN. To obtain ssDNA the distance between beads was gradually increased until the tension reached 70-80 pN, afterwhich the tether was incubated under this tension for 10-20 seconds. Typically, the slight increase in the end-to-end distance and a slight decrease in the tension were observed. After this, the DNA-tether was relaxed and its force-extension was measured. If the force extension curve followed that of ssDNA, then after incubation of DNA tether under low tension for about 7 seconds its force extension curve was measured once more to verify that dsDNA was fully melted into ssDNA.

DNA-tether formation and ssDNA generation using a longer DNA construct (22 and 24 kbp) was performed analogously except that a lower concentration of DNA fragments was used: 3.5 pM instead of 15 pM.

### Manipulations with RecA-ssDNA filament

#### RecA-ssDNA filament formation

The assembly of RecA-ssDNA filaments was performed by applying a stretching force of 12 pN to the ssDNA molecule in the channel containing 1 μM RecA and 1 mM ATP. The applied tension promoted the assembly of RecA filaments by removing the secondary structure of ssDNA. Binding of RecA to ssDNA resulted in the elongation of the DNA tether (Figure S5F).

#### The study of the effect of RecX on the active RecA-ssDNA filament

The effect of RecX on the active RecA-ssDNA filament was examined using the configuration of microfluidic channels presented in Figure S5B. After the assembly of the RecA-ssDNA filament in the channel supplemented with 1 μM RecA and 1 mM ATP, the applied tension was lowered to 3 pN. Under this tension the RecA-ssDNA filament was moved to the channel which in addition to 1 μM RecA and 1 mM ATP was also supplemented with RecX of indicated concentration and the change in the end-to-end distance was monitored.

To investigate the reversibility of the change in RecA-ssDNA filament length caused by RecX, another configuration of microfluidic channels was used (Figure S5C). The RecA-ssDNA filament was assembled in 2nd channel, afterwhich at 3 pN tension the length of the RecA-ssDNA filament was registered during incubation and transitions between 3rd and 4th channels, where the 3rd channel contained 1 mM ATP and the 4th channel contained 1 mM ATP and 500 nM RecX. Importantly, both 3rd and 4th channels contained no free RecA. The transfer of the filament between the midpoints of the adjacent channels took less than 5 seconds.

#### Manipulations with inactive RecA-ssDNA filament

The inactive RecA-ssDNA filament was obtained by transferring the preassembled active RecA-ssDNA filament to the ATP-lacking channel (Figure S5D). During transitions and incubations a constant tension of 3 pN was applied to the RecA-ssDNA filament. To examine the effect of RecX on the inactive RecA-ssDNA filament, it was transferred to the 4th channel supplemented with RecX of indicated concentration.

To examine the effect of RecX on the conformational transition from inactive state to the active, the inactive RecA-ssDNA filament was incubated in the presence of RecX (4th channel) at 3 pN tension and then was moved to the 2nd channel supplemented with 1 μM RecA and 1 mM ATP, where the increase in the RecA-ssDNA filament length was observed due to conformational transition induced by ATP binding. After the filament length was restored, the RecA-ssDNA filament was again moved to the 3rd channel, which caused a transition to an inactive state. In the 3rd channel the filament was incubated for 30 seconds, afterwhich the filament was moved to the 2nd channel, and dynamics of restoration of the RecA-ssDNA filament length was registered repeatedly.

#### Manipulations with RecA-dsDNA filament

To assemble RecA-dsDNA filament, dsDNA was incubated under 55 pN tension in the presence of 1 μM RecA and 1 mM ATP. After the assembly of the RecA-dsDNA filament, the applied tension was lowered to 3 pN for further manipulations. The measurements of the effect of RecX on the RecA-dsDNA filaments were performed similarly to the experiments with RecA-ssDNA filament.

#### RecX_mNG_ fluorescence intensity analysis

To calculate average intensity of the tether after incubation with fluorescent RecX_mNG_ a custom Fiji ImageJ script was used. First, mean intensity of the background was calculated by measuring mean gray value of the first 8 rows of pixels (8×52 pixels rectangle, 1 pixel = 142 nm) containing neither the tether nor the beads. This value was further subtracted from all the pixels in the image. To avoid the influence of auto-fluorescence of the beads only a central part of the tether comprising a rectangle of 21×6 pixels (for apo state) and 31×6 pixels (for ATP state) were used to calculate an average intensity of the fluorescent signal.

## Acknowledgements

We are grateful to Elena Znobishcheva (Peter the Great St Petersburg Polytechnic University, St Petersburg) for her help in the preparation of the DNA construct used for fluorescence experiments.

This research was funded by the Russian Science Foundation, grant number [19–74-10049].

---

Supplementary figures–source data 1

Source data for plots in figures S2, S3, S4, S5F

**Figure S1.**
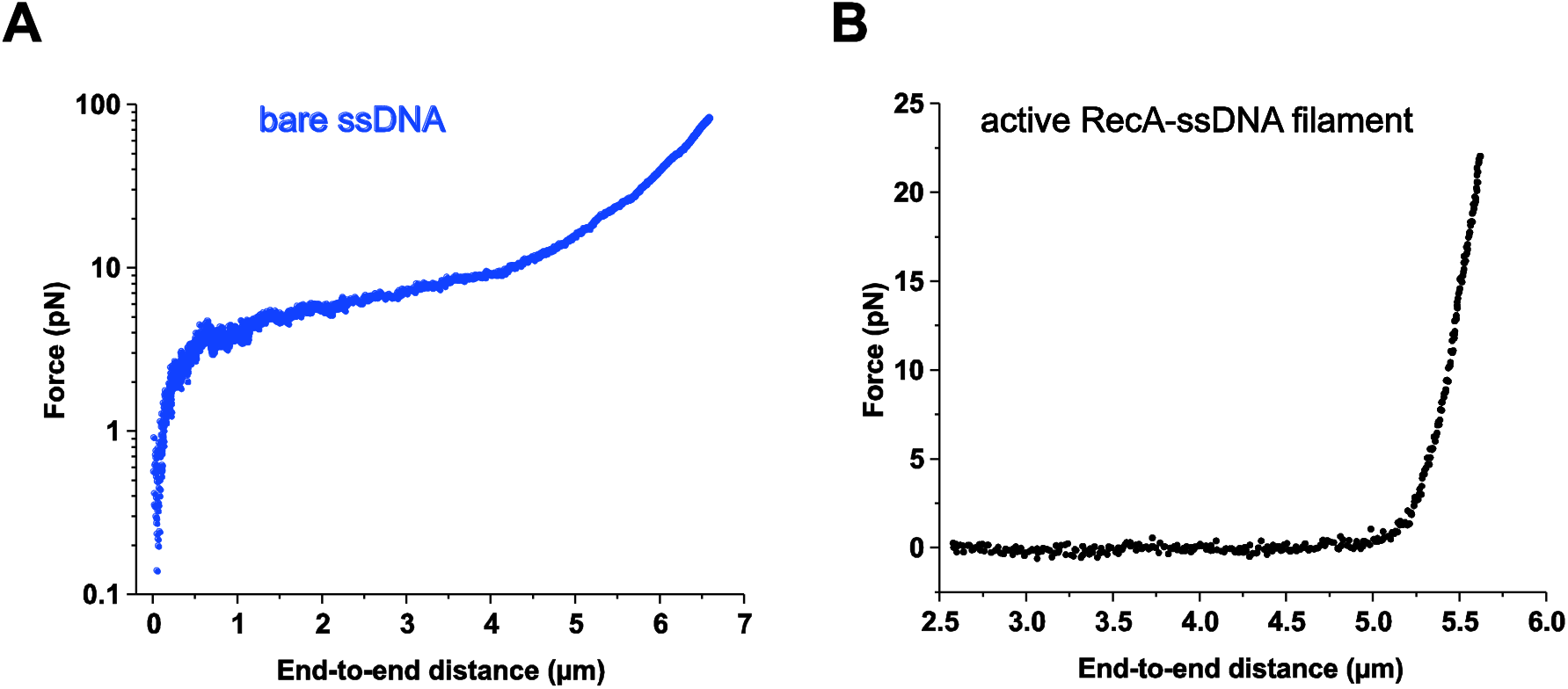
The comparison of force-extension behaviour of bare ssDNA (a) and the ATP-bound RecA-ssDNA filament (b). Adopted from [1].

**Figure S2.**
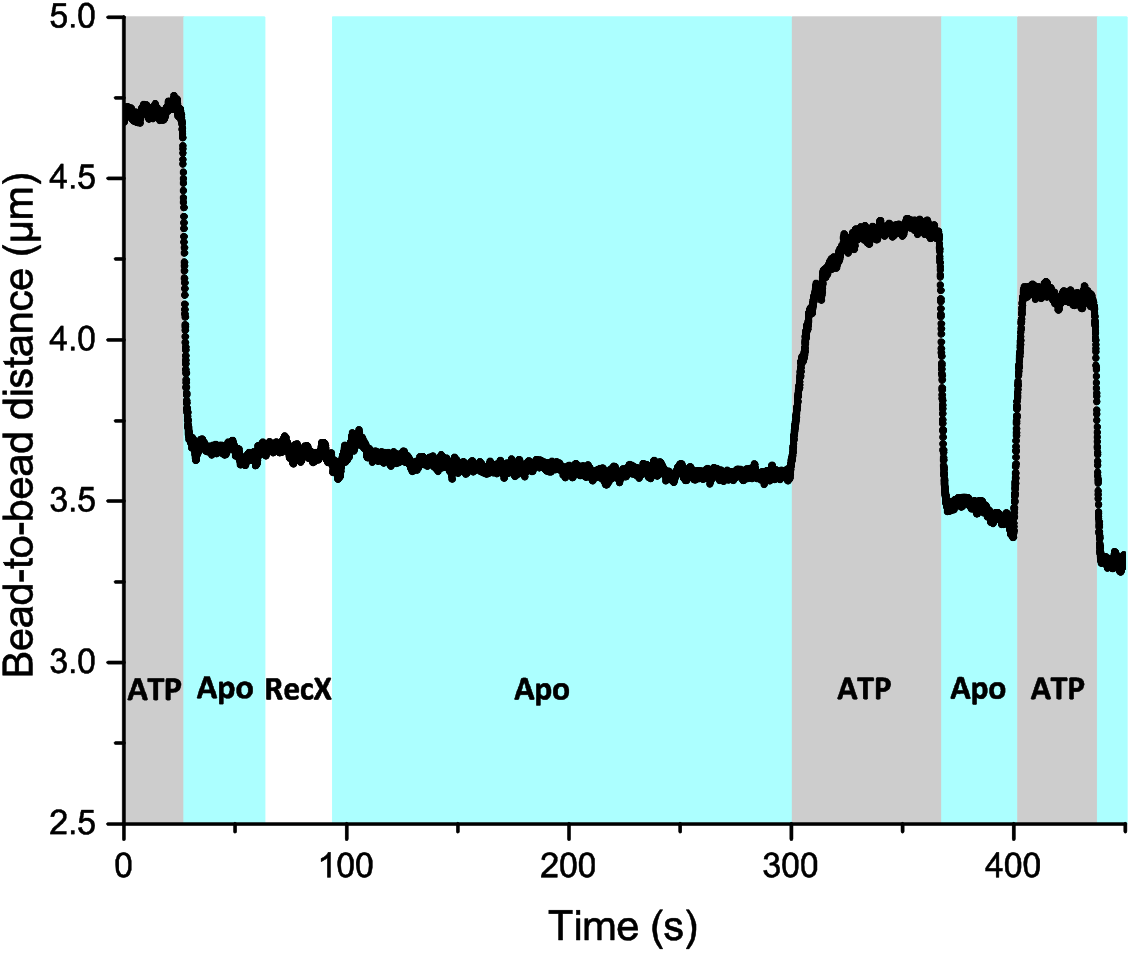
The effect of the slowed-down decompression retains when RecA-ssDNA filament is incubated in the RecX-free buffer after short incubation with RecX. ATP-containing channel was also supplemented with free RecA.

**Figure S3.**
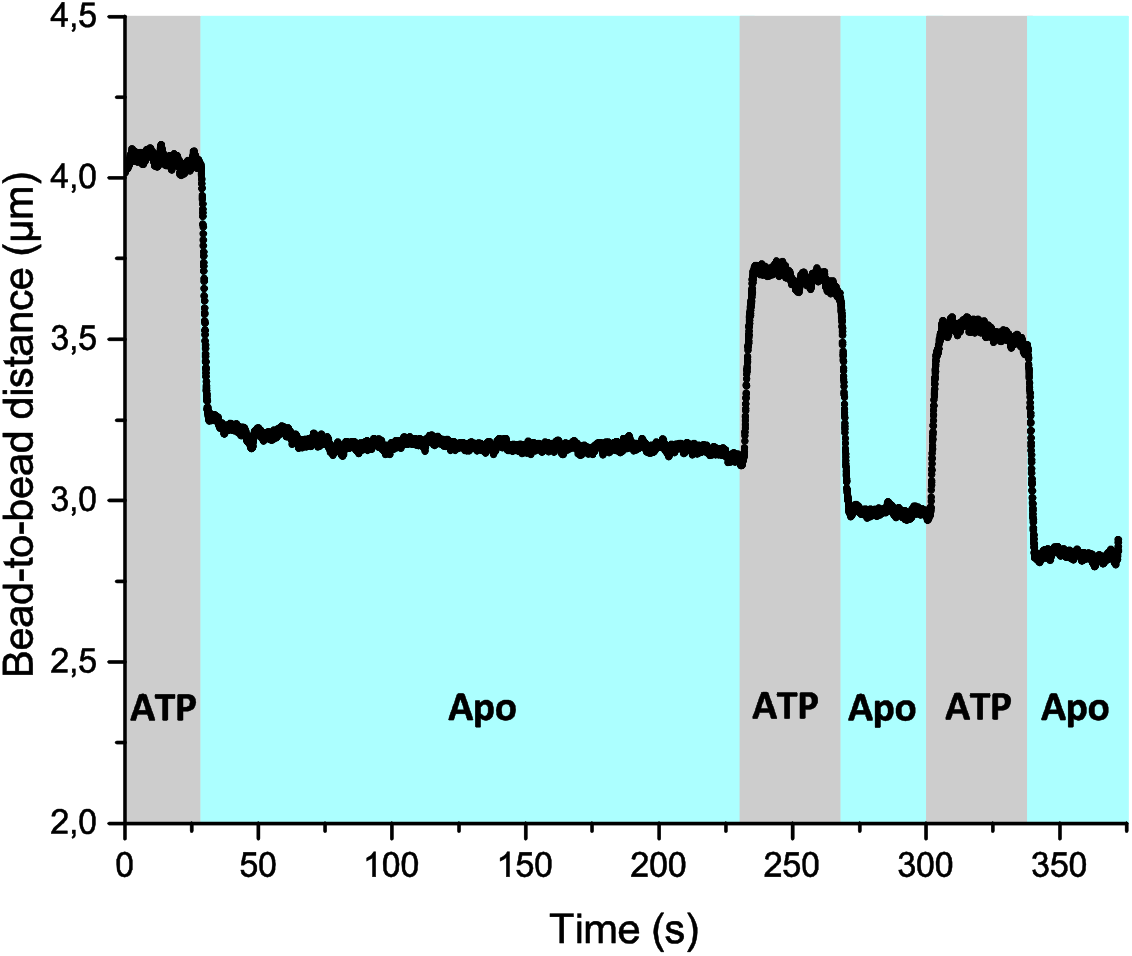
The effect of the slowed-down decompression is independent of incubation time of the RecA-ssDNA filament in the Apo channel in the absence of RecX. ATP-containing channel was also supplemented with free RecA.

**Figure S4.**
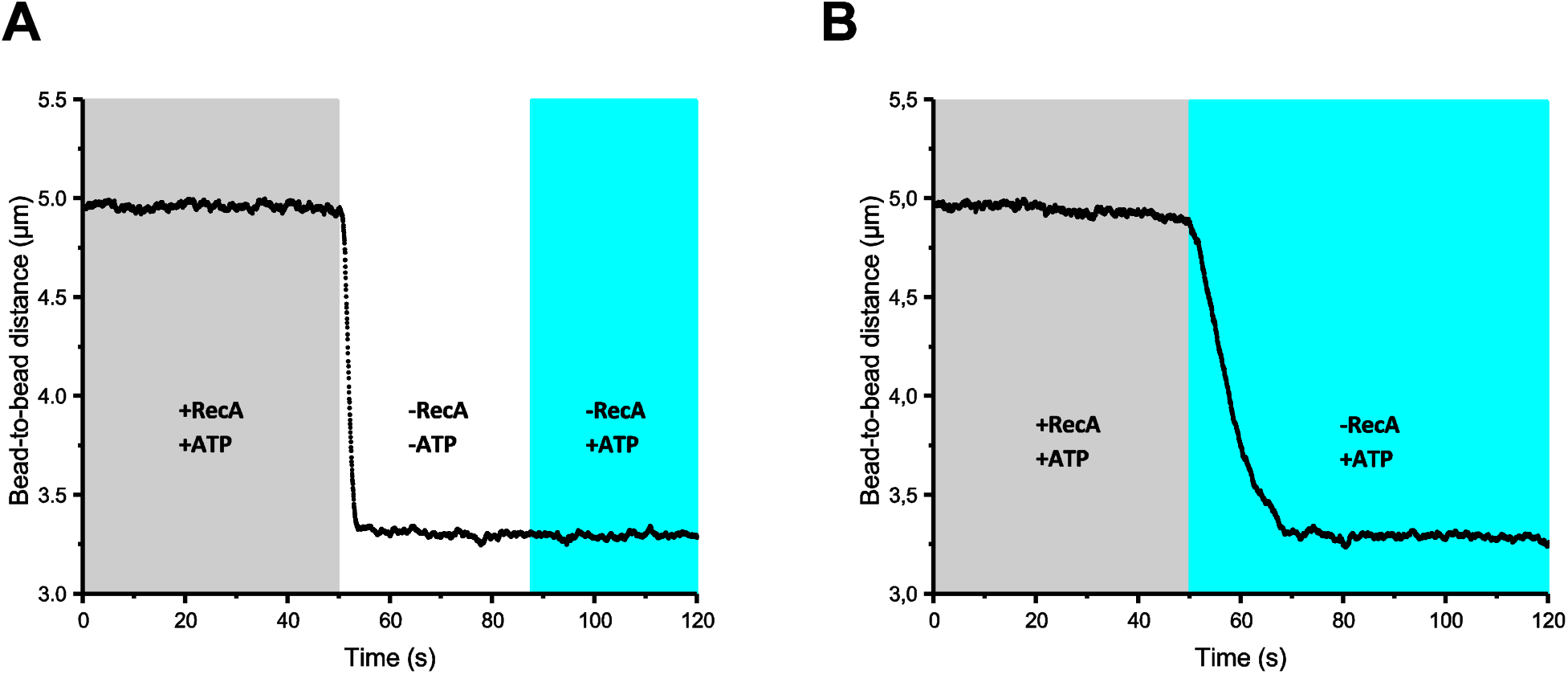
RecA-dsDNA filament is stable only in the presence of both free RecA and ATP. (a) ATP elimination leads to rapid RecA-dsDNA filament disassembly. (b) The elimination of free RecA promotes RecA-dsDNA filament disassembly.

**Figure S5.**
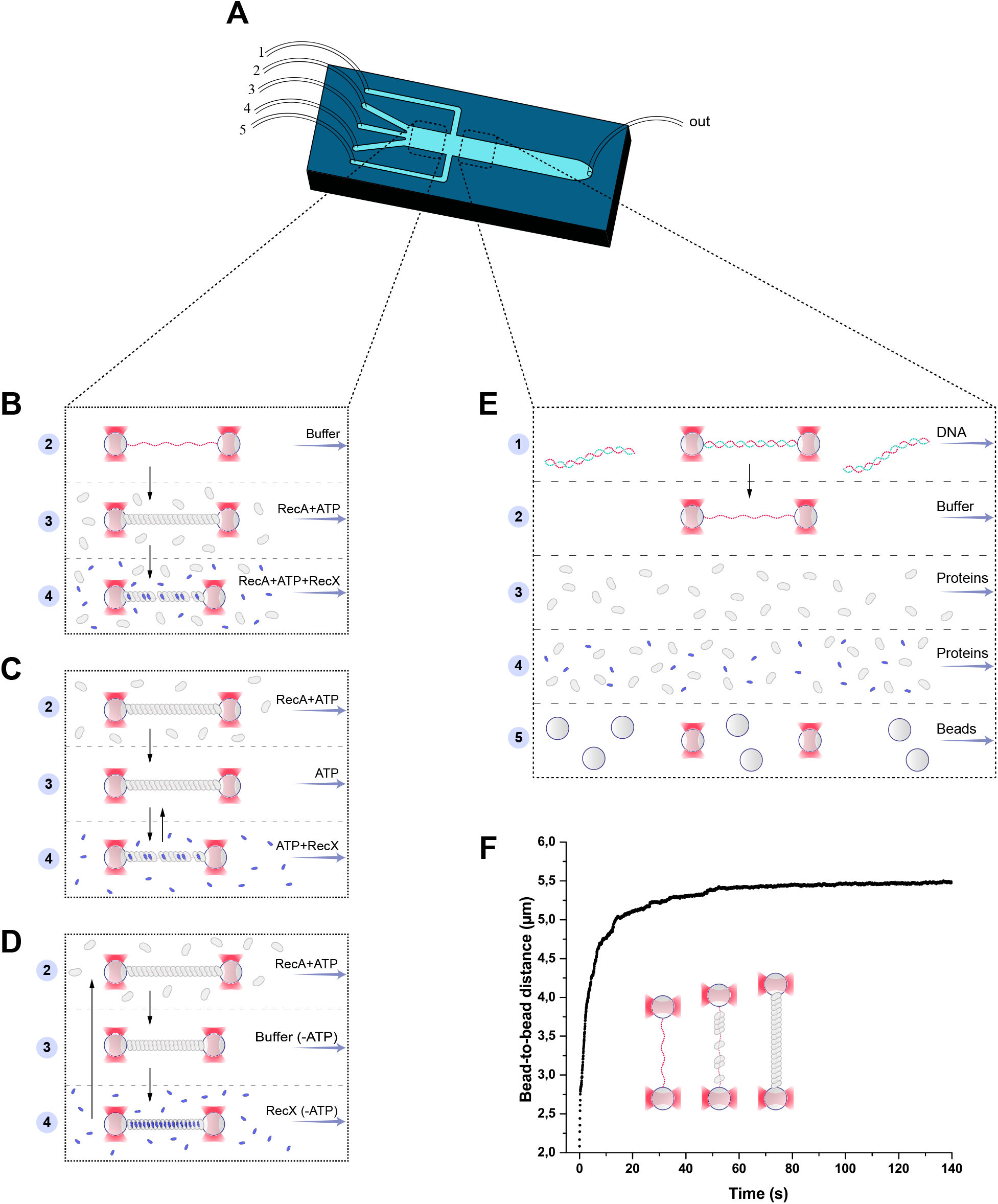
Single-molecule assay. Single-molecule manipulations were performed within a 5-channel microfluidic flow chip (a). Two working regions are highlighted with a dash line. The three-channel region (left) was used to study the effect of RecX on the RecA-ssDNA filament (b-d). In the 5-channel region the beads trapping, DNA-tether formation and generation of ssDNA by force-induced melting were performed (e). The RecA-ssDNA filaments were assembled by applying a stretching force of 12 pN to the ssDNA molecule in the channel containing 1 μM RecA and 1 mM ATP. Binding of RecA to ssDNA was followed by an increase in the end-to-end distance (f).

## Notes

### Competing Interest Statement

The authors have declared no competing interest.

